# Exploiting evolutionary trade-offs to combat antibiotic resistance

**DOI:** 10.1101/2020.01.20.912303

**Authors:** Sergey Melnikov, David L. Stephens, Xian Fu, Hui Si Kwok, Jin-Tao Zhang, Yue Shen, Jeffery Sabina, Kevin Lee, Harry Lee, Dieter Söll

## Abstract

Antibiotic resistance frequently evolves through fitness trade-offs in which the genetic alterations that confer resistance to a drug can also cause growth defects in resistant cells. Here, through experimental evolution in a microfluidics-based turbidostat, we demonstrate that antibiotic-resistant cells can be efficiently inhibited by amplifying the fitness costs associated with drug-resistance evolution. Using tavaborole-resistant *E. coli* as a model, we show that genetic mutations in leucyl-tRNA synthetase (that underlie tavaborole resistance) make resistant cells intolerant to norvaline, a chemical analog of leucine that is mistakenly used by tavaborole-resistant cells for protein synthesis. We then show that tavaborole-sensitive cells quickly outcompete tavaborole-resistant cells in the presence of norvaline due to the amplified cost of the molecular defect of tavaborole resistance. This finding illustrates a potentially generalizable approach for combating therapeutic resistance, prolonging the effectiveness of drugs and enabling the use of drugs that are no longer effective due to the rapid evolution of resistance.

---

The therapeutic use of drugs against microbial pathogens and cancer is currently undergoing a paradigm shift from *traditional therapies* toward *adaptive therapies*, in which disease is treated as an evolutionary process to minimize the risk of drug resistance evolution^1-5^.

With traditional therapy, the goal is to eradicate the disease by eliminating every pathogenic cell in the human body via the long-term administration of a drug at the maximum tolerable dose. A notorious drawback of this approach is that, after an initially effective treatment stage, it frequently results in the development of drug resistance during later treatment stages. In contrast, adaptive therapies aim to manage the disease without necessarily eradicating it. Evidence from studies of adaptive therapies suggest that – rather than forcing pathogenic cells into evolving elaborate forms of drug resistance during the long-term administration of antimicrobial or anti-cancer drugs – pathogens should be treated using repeated, short-term bouts of drug application that are interrupted by periods without treatment^3,5^. The rationale for this approach is that, during these periods without treatment, resistant cells will be naturally outcompeted by non-resistant cells (or cells with lower levels of resistance) either due to genetic drifts or because the resistant cells are frequently less fit than non-resistant cells in the absence of drug^1,2^.

Multiple studies have shown that integrating this evolutionary principle into clinical treatment protocols can drastically improve disease progression in patients suffering from advanced forms of infectious disease or cancer^1,3-6^. For instance, adopting this approach dramatically improved the effectiveness of abiraterone-mediated treatment of metastatic forms of prostate cancer, where just one patient out of 11 developed therapeutic resistance during a course of adaptive therapy, compared with 14 out of 16 patients who developed therapeutic resistance following traditional therapy^5^. Therefore, it is becoming increasingly important to learn whether we can reduce the periods without treatments in order to combat therapeutic resistance by encouraging natural competition between resistant and non-resistant cells.

In this study, we sought to exploit the evolutionary trade-off associated with drug resistance evolution as a means to promote competition between resistant and non-resistant cells. Our aim was to capitalize on the fact that drug resistance frequently evolves as a trade-off in which the very same genetic mutations that alter a drug target and decrease the affinity of a target for the drug, can also compromise the biological activity of the target. This compromised activity leads to growth defects in drug-resistant cells in the absence of drug – a scenario that is especially common for synthetic drugs that act on a single target in the cell^7-12^. For instance, numerous studies showed that evolution of drug resistance in bacterial and cancer cells can lead to collateral sensitivity (a phenomenon in which resistance to one drug makes cells more sensitive to another drug), making it possible to exploit these evolutionary trade-offs to control drug resistance evolution^13-17^.

In this study, we tested our hypothesis that drug-resistant cells can be selectively inhibited by enhancing toxicities associated with drug resistance, i.e. by placing them under conditions in which the molecular defects associated with drug resistance are especially fitness-costly.

For our drug model, we used tavaborole (also known as AN2690), a synthetic small-molecule inhibitor of protein synthesis^18-20^. In 2014, this drug was approved by the FDA to treat onychomycosis (nail fungus)^21^. Numerous studies have indicated that tavaborole and its chemical derivatives may also be of potential use against microbial pathogens, including *Plasmodium falciparum*^22,23^, *Toxoplasma gondii*^24^, *Trypanosoma brucei*^25-29^, *Cryptosporidium parvum*^24^, *Staphylococcus aureus*^30^, *Mycobacterium tuberculosis*^31-34^, *Streptococcus pneumoniae*^35^, and multidrug-resistant strains of *Pseudomonas aeruginosa* and *Escherichia coli*^20^.

Tavaborole inhibits cell growth by inactivating leucyl-tRNA synthetase (LeuRS), an essential enzyme that attaches leucine to its corresponding tRNA (tRNA^Leu^) to produce leucyl-tRNA^Leu^, a substrate used in protein synthesis. LeuRS has two catalytic centers: the synthesis site, which is responsible for leucyl-tRNA^Leu^ formation, and the editing site, which is responsible for quality control during leucyl-tRNA^Leu^ formation. When the synthesis site makes occasional errors by attaching amino acids other than leucine to tRNA^Leu^, the editing site detaches these amino acids from tRNA^Leu^, thereby preventing errors in protein synthesis^36^. Tavaborole inhibits LeuRS by targeting its editing domain, where tavaborole covalently binds with tRNA^Leu^ and prevents the dissociation of tRNA^Leu^ from LeuRS, leading to the inhibition of both protein synthesis and cell growth^19^.

Previous studies in the model organism *Saccharomyces cerevisiae* have suggested that resistance to tavaborole can evolve via mutations in the *leuS* gene (the gene that codes for leucyl-tRNA synthetase): these mutations occur in the editing domain of LeuRS and frequently impair the protein’s editing activity^19,37-39^. A similar resistance mechanism was observed in clinical isolates of tavaborole-treated multidrug-resistance *Escherichia coli*^40^, and in *Staphylococcus aureus* that were treated with a tavaborole derivative, AN3365, the phase II clinical trials of which have been suspended due to the rapid development of AN3365 resistance^39^. In the present study, therefore, we endeavored to determine if it is feasible to use this development of resistance, which likely arises at the expense of LeuRS editing, as a weakness via which tavaborole-resistant cells could be inhibited.

We first tested how frequently tavaborole resistance originates from mutations in LeuRS. For this purpose, we evolved tavaborole-resistant *E. coli*. We used six initially identical populations of *E. coli*. We cultured these populations over 8 days, with the daily transfer of 5% of each culture into fresh media supplemented with tavaborole, gradually increasing the tavaborole concentration from sub-MIC (3 µg/ml) to doses above the MIC (160 µg/ml) (**Materials and methods, Fig. S1**). By day 8, all populations were able to rapidly grow in the presence of tavaborole, indicating the development of tavaborole resistance (**Fig. S1**).

We next tested whether the resistant cells had mutations in the *leuS* gene. First, we sequenced the *leuS* gene in 120 tavaborole-resistant colonies (20 colonies selected at random from each of the six evolved populations) and found that one or two mutations in *leuS* were present in organisms from each colony. All these mutations were located in the *leuS* segment corresponding to the editing domain of LeuRS (**Fig. 1A, Table S1, Supplementary Data 1**). We next tested whether it was these mutations (and not other mutations possibly present in the *E. coli* genome) that conferred tavaborole resistance. We selected five of the tavaborole-resistant colonies that contained one of the five most frequently observed mutations, including G225E, G229V, Y330F, G331S, and R344S. In each of these colonies, we replaced the mutated *leuS* gene with the wild-type *leuS* gene and found that all of the derived clones had lost their tavaborole resistance, indicating that tavaborole resistance was conferred by mutations in the *leuS* gene (and not by other mutations that may possibly be present elsewhere in the genome) (**Fig. S2A**). In a complementary experiment, we used wild-type *E. coli* and mutated their *leuS* gene by introducing one of the five frequently observed LeuRS mutations, G225E, G229V, Y330F, G331S, or R344S. We found that each of these mutations conferred resistance to tavaborole (10 µg/mL) upon their insertion into the wild-type *E. coli* genome, illustrating that each of these mutations on its own is sufficient to endow *E. coli* with a high level of resistance to tavaborole (**Fig. S2B**). Finally, we performed time-resolved whole-population genome sequencing of the two evolving populations (lineages A and B) and found that mutations in the *leuS* gene were the only detectable mutations whose frequency gradually increased during the course of the experiment and which were present in the majority of cells by the end of the experiment (**Fig. S3, Supplementary Data 3**). Taken together, our data showed that – not only *leuS* mutations may confer resistance to tavaborole, as previously observed^19,37-39^ – but, at least in our experimental conditions, they appear to be the most preferential route for tavaborole resistance evolution.

**Figure 1.**
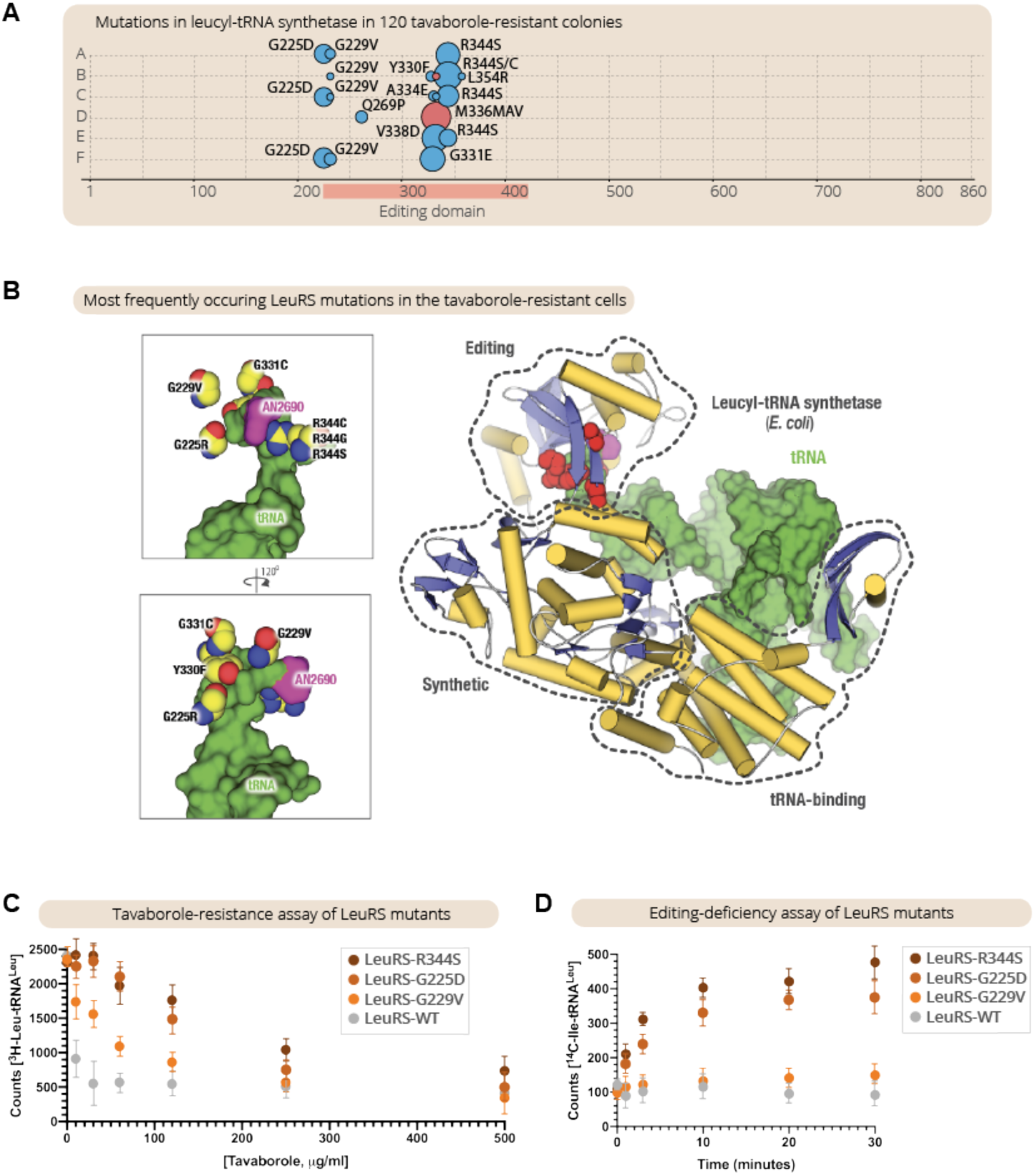
Impairing LeuRS editing activity is the primary route of tavaborole-resistance evolution in continuous *E. coli* cultures. **A.** The panel summarizes mutations in the *leuS* gene that were observed in 120 tavaborole-resistant colonies (20 colonies from each of the six tavaborole-resistant lineages of *E. coli*). The size of the circles indicates the frequency of the observed mutations, including amino acid substitutions (blue circles) and repeat insertions (red circles). As the panel shows, all the observed mutations were located in the editing domain segment of the *leuS* gene, with one or two mutations per colony. **B.** The crystal structure of LeuRS in complex with tRNA^Leu^ and tavaborole shows the location of tavaborole-resistance mutations in LeuRS synthetase. Remarkably, all the observed mutations are located not at the interface between LeuRS and tavaborole but at the interface between LeuRS and tRNA^Leu^, suggesting these mutations prevent tavaborole inhibition via impaired editing activity. **C.** A tavaborole-resistance assay shows IC_50_ measurements, in which LeuRS activity (assessed by monitoring leucyl-tRNA^Leu^ formation) was measured in the presence of tavaborole (0–500 µg/mL). **D.** An editing activity assay shows that, unlike the wild-type LeuRS, the LeuRS-resistant mutants can erroneously charge tRNA^Leu^ with isoleucine, leading to the formation of an erroneous reaction product, isoleucyl-tRNA^Leu^ (Ile-tRNA^Leu^). Collectively, the figure illustrates that – not only tavaborole resistance may sometimes evolve through mutations in the editing domain of LeuRS, as previously observed in individual clones^19,37-39^ – but these mutations represent the primary route of the resistance evolution, with the most frequently observed mutants being the most editing-defective.

We next tested whether the tavaborole-resistance mutations affected the editing activity of LeuRS. First, we mapped the observed mutations onto the previously determined structure of LeuRS synthetase bound to tavaborole-modified tRNA^Leu^ (**Fig. 1B**). Remarkably, none of the mutations were located at the interface between LeuRS and tavaborole (**Fig. 1B**). Instead, the mutations were clustered at the interface between tRNA^Leu^ and LeuRS, suggesting that tavaborole resistance is likely acquired not via the direct prevention of drug binding but because LeuRS mutants cannot properly bind tRNA^Leu^ within the editing site. This could be due either to steric clashes between tRNA^Leu^ and LeuRS (as seems to be the case for G225E, G229V, and G331C mutants) or to disrupted tRNA-LeuRS contact (as seems to be the case for R344C/S and Y330F mutants) (**Fig. 1B, Fig. S4**). We then purified LeuRS mutants carrying the most frequently observed mutations (G225E, G229V, and R344S) and tested their activity *in vitro* by measuring kinetics of tRNA^Leu^ aminoacylation with leucine (to assess the synthetic activity of LeuRS) or isoleucine (to assess the editing activity of LeuRS). Consistent with the structural observations, the *in vitro* measurements showed that all mutants tested exhibited high levels of resistance to tavaborole (**Fig. 1C**), but at the same time were also editing-defective (**Fig. 1D**). Thus, we found that – not only some of the tavaborole-resistance mutations may impair the editing activity of LeuRS, as observed previously in individual resistant clones^19,37,38^ – but the most frequently-occurring mutations are the ones that impair the editing activity most, with the majority of resistant cells in the evolving populations being editing-defective.

Our final investigation sought to determine if we could exploit these editing defects in LeuRS to inhibit tavaborole-resistant cells. Previously, laboratory-engineered *E. coli* strains with editing-deficient LeuRS were shown to be hypersensitive to the toxicity of chemical analogs of leucine (including norleucine, norvaline, homocysteine, and homoserine); in these strains, and in some naturally occurring parasites^41,42^, the absence of editing activity caused leucine analogs incorporation into protein sequences, resulting in protein misfolding and cell growth arrest^43,44^ as a result of mistranslation^45^. We therefore anticipated that tavaborole-resistant cells would also be hypersensitive to the toxicity of leucine analogs.

To test this hypothesis, we used a competition assay in which we mixed tavaborole-resistant cells with wild-type *E. coli* expressing GFP, at a ratio of approximately 1:1, and incubated the cell mixture in a microturbidostat for 5 days (**Fig. S5**)*. We observed that, in the absence of leucine analogs, the ratio between the two populations remained largely unaltered throughout the experiment, indicating comparable growth rates of both the resistant and non-resistant *E. coli* populations (**Fig. 2**). However, when the growth media was supplemented with norvaline (0.5 mM), the tavaborole-resistant population rapidly declined to less than 3% of the total cell count (**Fig. 2**). Thus, this experiment illustrated that tavaborole-resistant *E. coli* are indeed vulnerable to the toxicity of leucine-like amino acids, showing that we can capitalize on the genetic defects of tavaborole-resistant populations to create the additional burden (negative selective pressure) associated with antibiotic-resistance mutations and selectively inhibit antibiotic-resistant cells.

**Figure 2.**
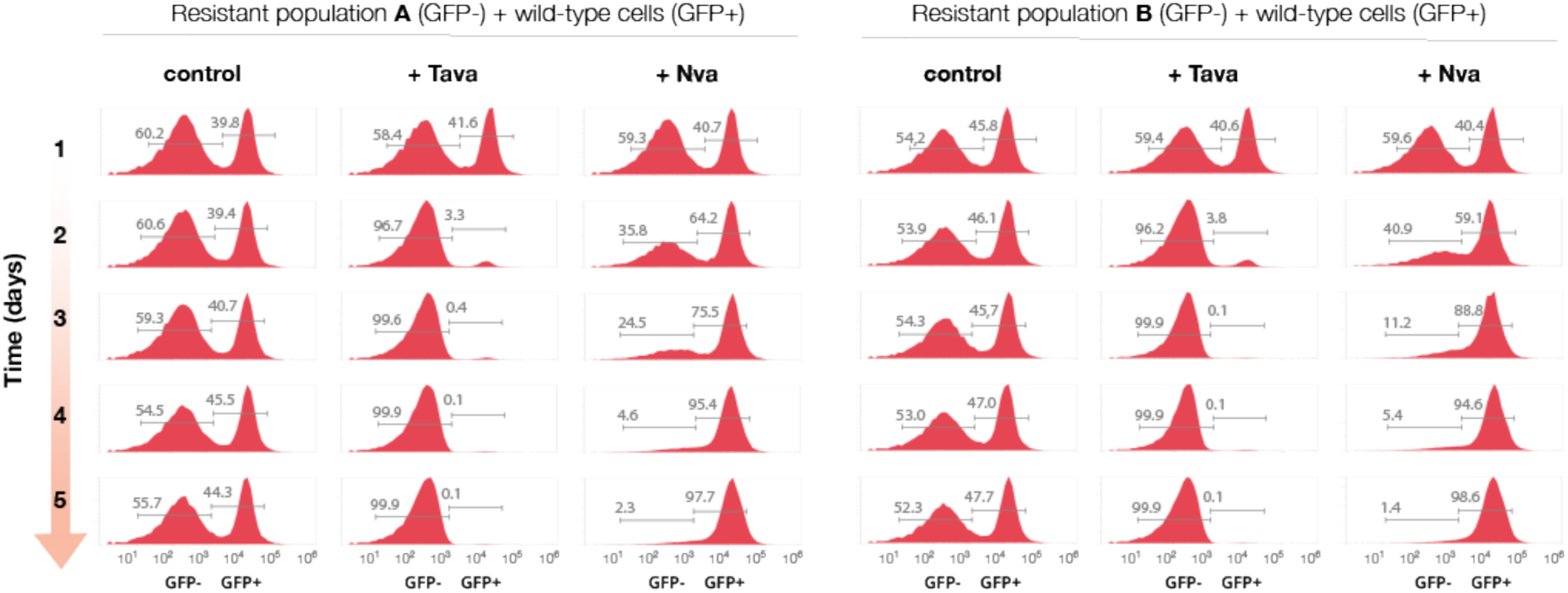
Tavaborole-resistant cells can be rapidly and selectively suppressed under conditions that require LeuRS editing activity. Flow-cytometry analysis of a competition assay in which two populations of *E. coli* were mixed at a ratio of approximately 1:1 and grown for five days either in chemically defined media (“control”), or in the same media supplemented with tavaborole (“+Tava”, 1 µg/mL) or norvaline (“+Nva”, 0.5 mM). The competing populations were (i) wild-type *E. coli* that expressed genome-encoded GFP (corresponding to the GFP-positive peaks) and (ii) tavaborole-resistant *E. coli* (population A or population B corresponding to the GFP-negative peaks).

Overall, our case study illustrates that the natural competition between resistant and non-resistant cells can be drastically enhanced by exploiting the burden associated with those mutations that confer antibiotic resistance. In the case of tavaborole, the fitness cost associated with LeuRS editing deficiency can be amplified by adding norvaline to the growth media, making LeuRS editing defects highly disadvantageous for *E. coli* growth and leading to the rapid and selective suppression of drug-resistant *E. coli* populations.

Given that resistance to many synthetic drugs evolves primarily through alterations of a drug’s target^11^, it is entirely possible that a conceptually similar approach could be applied to develop anti-resistance strategies for other drugs. By encouraging the natural competition between resistant and non-resistant cells during the “off”-phases of the “on-off” cycles of adaptive therapy (e.g. by administration of norvaline following the cycle of tavaborole administration), we can potentially accelerate adaptive therapies by significantly shortening their drug-free phases. Thus, our study provides an important opportunity and adds to the current arsenal of strategies (including collateral sensitivity, spatial competition constrains^46^, containment strategy^47^, and others^48^) of how to use a drug to minimize the threat of drug resistance evolution.

## MATERIALS AND METHODS

### Evolution of tavaborole resistance in *E. coli*

The tavaborole-resistant *E. coli* were evolved by growing six initially identical populations of BL21(DE3) in LB media at 37°C using 12-well plates (Cornig, Ref. 353047) that were incubated in Synergy|HTX multi-mode plate reader (BioTek). Each population was growing in 1.5 ml of LB media, and, each day, 5% of each of the evolving cell cultures were transferred into a new plate where each culture was diluted with 1.5 mL of fresh LB medium. During the 8 days of the experiment, the tavaborole concentration was gradually increased from the initial of 2.5 µg/mL to the final of 180 µg/mL (**Fig. S1**). The cell growth during the experiment was continuously assessed by making one OD_600_ measurement per minute for each of the evolving cultures.

### Sequencing of *leuS* gene in tavaborole-resistant *E. coli* colonies

To determine the sequence of *leuS* gene in tavaborole-resistant cells, each of the evolved *E. coli* populations was plated on LB-agar plate supplemented with tavaborole (60 µg/mL), grown overnight at 37°C. Then, and then 20 individual colonies were picked at random from each of the six plates. Each of these colonies was then used as a DNA template for PCR amplification of *leuS* sequence. To amplify *leuS* gene, we used Phusion® High-Fidelity DNA Polymerase (NEB) and primers **1** and **2** (here and below we refer to the primers according to the **Table S2, Supplementary Data 1**). The amplified gene was analyzed by Sanger sequencing (The Keck DNA Sequencing Facility at Yale School of Medicine) using each of the following primers for sequencing: **1, 3**, and **4**.

### *E. coli* genetics to alter *leuS* sequence

To introduce tavaborole-resistance mutations in *leuS* gene in *E. coli* genome, we used the previously published protocol for recombineering, a homologous recombination-based method of genetic engineering^49^. In brief, the editing domain-coding segment of *leuS* gene was PCR amplified by using Phusion® High-Fidelity DNA Polymerase (NEB), primers **1** and **2**, in and the tavaborole-resistant colonies as DNA template. The PCR product was used for transformation of the electro-competent *E. coli* (BL21(DE3)) that were provided with λ Red recombination genes by using pSIM6 plasmid, as described in “Provision of the λ Red recombination genes”^49^. The transformed cells were then grown for 1h in LB medium at 37°C, transferred onto LB-agar plates supplemented with tavaborole (10 µg/mL) and grown overnight at 37°C. The resulting clones were grown in LB media, analyzed by Sanger sequencing of PCR-amplified *leuS* gene (using primers **3** and **4**) and used for growth assessment at 37°C in LB media by using Synergy|HTX multi-mode plate reader (BioTek).

To replace the mutated *leuS* gene in tavaborole-resistance cells with the wild-type *leuS*, we fused PCR-amplified *leuS* gene (using primers **3** and **4** and an aliquot of the wild-type *E. coli* as a DNA template) with the *Neo*-selection gene by using In-Fusion HD Cloning kit (Takara), and integrated the resulting *leuS-Neo* DNA into genomes of the tavaborole-resistant cells by using the scarred method for insertion of a nonselectable DNA fragment recombineering protocol with *Neo* gene as a selectable marker^49^.

### Library preparation and whole genome sequencing

Genomic DNA from the evolving *E. coli* populations was purified by using 1 mL of each *E. coli* sample that was treated by using QIAamp DNA Mini Kit (Quiagen). For the sequencing library construction, the genomic DNA was fragmented by ultrasonication to the size between 50 bp and 800 bp. The DNA fragments (from 100 bp tod 300 bp) were selected and further capped with the flanking dATP ends. The dTTP tailed adapters were ligated to both ends of the DNA fragments. The ligation product was then amplified and subjected to the single-strand circularization process. The remaining linear molecule was digested with the exonuclease to generate a single-strand circular DNA library. Sequencing was conducted according to the standard BGISEQ-500 protocol. Zebracall (a base calling software) was used to make raw reads.

### Sequencing quality control and mapping

Quality control of sequencing reads was performed before mapping. Reads with adapters and shorter than 100 bp were removed. Reads containing more than 1% of unknown base and containing more than one base with Phred-score lower than 10 were removed. The quality control step resulted in cleaned paired-end reads with more than 1000-fold sequencing depth of the reference genome. The reads after filtration were mapped to reference sequences using BWA 0.5.6 according to the standard settings^50^. For each alignment result, local realignment was performed with GATK 2.7 built-in RealignerTargetCreator and IndelRealigner tools to clean up mapping artifacts due to reads mapping on the edges of indels^51^. The resulting files in BAM format were then prepared for initial SNV/indel calling.

### Identification of SNVs and indels, and data deposition

GATK 2.7 pipeline^51^ was utilized to identify the SNVs and indels using default parameters. The variants were filtered by the following criteria: QUAL<50 or FS>3 or BaseQRankSum>3 or MQRankSum>3 or ReadPosRankSum>3 or MQ<10 or DP<10. Next, annotation was performed for observed variants types including synonymous type, nonsynonymous type, frameshifts, and variant outside the coding region. The sequencing data were then deposited to the CNGB Nucleotide Sequence Archive (CNSA: https://db.cngb.org/cnsa; accession number CNP0000842) (the links to the datasets are provided in the **Supplementary Data 2**).

### Preparation of tRNA^Leu^ transcript for biochemical assays

To produce tRNA^Leu^ transcript for enzymatic assays, we used the tRNA^Leu^ sequence for tRNA^Leu^CAA-1-1 from *E. coli* K-12 MG1655 strain (http://gtrnadb.ucsc.edu/genomes/bacteria/Esch_coli_K_12_MG1655/genes/tRNA-Leu-CAA-1-1.html). We first produced the DNA-template for T7-transcription using PCR with Taq polymerase (NEB) and primers **11, 12**, and **13**. We then used HiScribe™ T7 High Yield RNA Synthesis Kit (NEB) to produce tRNA^Leu^. For this purpose, 4 µg of the DNA template were added to the reaction mixture (the total volume of 40 µL), containing 10 mM ATP, 10 mM CTP, 10 mM GTP, 10 mM UTP, 200 mM of GMP (to prevent the presence of phosphates at the tRNA^Leu^ 5’-end) and 4 µL of T7 RNA polymerase solution. The reaction mixture was incubated at 40°C for 15 hours, treated with 2 units of DNase I (NEB) at 37°C for 1 hour followed by DNase I heat inactivation. The resulting mixture was diluted to 100 µL with water, mixed with 100 µL of acid phenol (pH 4.5):chloroform:IAA (125:24:1). After the extraction, the aqueous phase was used for size-exclusion chromatography by using a disposable PD SpinTrap G-25 column (GE Healthcare Life Sciences). The purified tRNA^Leu^ was then ethanol precipitated, dissolved in water, analyzed by 12% PAAG/7M urea, and stored at 2 µg/µL concentration at −20°C.

### Purification of LeuRS and its mutants for biochemical assays

To purify recombinant LeuRS synthetase, *leuS* gene and its mutants were PCR-amplified by using primers **3** and **4** and genomic DNA from either wild-type or tavaborole-resistant *E. coli* colonies as the template. The PCR products were cloned into pBAD28 vector (ATCC® 87400™) by using In-Fusion HD Cloning kit (Takara) by using primers **7** and **8** to amplify pBAD28 vector (**Table S2, Supplementary Data 3**). To express LeuRS, *E. coli BL21*(DE3) cells were transformed with LeuRS-coding plasmids, and the synthesis of recombinant proteins was induced at 16 °C by the addition of arabinose (to the final concentration of 0.2% w/v) to the growth medium. The proteins were then purified by the kit of Ni-affinity purification (BioRad), followed by ammonium sulfate protein fractionation and size-exclusion chromatography on a Superdex Increase 200 10/300 GL column (Pharmacia Biotech) in the aminoacylation buffer containing 60 mM Tris⋅HCl pH 7.5, 10 mM MgCl_2_, and 1 mM DTT. Then, the proteins were concentrated to 10 mg/mL using Amicon Ultracel centrifugal filters (molecular weight cutoff, 30 kDa) and immediately used in biochemical assays.

### LeuRS aminoacylation activity assay

To assess LeuRS aminoacylation activity, we used the previously described protocol monitoring synthesis of [^14^C]-labeled leucyl-tRNA^Leu 52^. In brief, we measured LeuRS activity in the aminoacylation buffer supplemented with 8 µM tRNA^Leu^ transcript, 40 µM L-leucine, 2 µM L-[^3^H]-leucine (PerkinElmer, 144 Ci/mmol), tavaborole (0-500 µg/mL) and a recombinant LeuRS from *E. coli* (the final concentration 20 nM). The reaction was initiated by the addition of LeuRS and terminated by the addition of trichloroacetic acid (TCA) to 5% v/v concentration. In the meantime, we prepared Whatman 3MM filter pads by adding 100 µL of 5% TCA onto each pad and drying each pad before the measurements. Then, 10 µL of each reaction mixture was transferred to the filter, and the filters were immediately soaked in 15 mL of 5% ice-cold TCA. Each filter was washed 3×20 minutes by using 15 ml of ice-cold 5% TCA to remove the free amino acid. Then each filter was washed by using 15 ml of 96% ethanol, dried, mixed with 5 mL of scintillation fluid (MP Biomedicals) and used for measurements by using LS 6500 scintillation counter (Beckman Coulter).

### LeuRS editing activity assay

To assess LeuRS editing activity, we measured the ability of LeuRS and its mutants to produce the misaaminocylated tRNA^Leu^, Ile-tRNA^Leu^, as described previously^53^. In brief, we measured LeuRS activity in the aminoacylation buffer supplemented with 8 µM tRNA^Leu^ transcript, 40 mM L-isoleucine (99%, W527602; Sigma-Aldrich), 2 µM L-[^14^C]-isoleucine (PerkinElmer, 322 mCi/mmol), tavaborole (0-500 µg/mL) and a recombinant LeuRS from *E. coli* (final concentration is 2 µM). The reaction was initiated by the addition of LeuRS and terminated by the addition of trichloroacetic acid (TCA) to 5% concentration. The amount of Ile-tRNA^Leu^ was then assessed by using aliquot transfer on Whatman 3MM filter pads, as described for the aminoacylation activity assay.

### Construction of GFP-encoding *E. coli*

The electrocompetent *E. coli* cells (BL21(DE3)) were transformed with the pOSIP-CT (TetR, P21) integration plasmid coding for sfGFP (**Supplementary Data 4**). The transformed cells were grown at 37°C for 1 hours in LB media and then for 12 hours in LB media supplemented with tetracycline (50 µg/mL). Then, the GFP-positive cells were sorted by using BD FACS Aria III Cell Sorter (BD Biosciences). The sorted cells were plated on a 100mm LB-agar Petri dish and grown at 37°C for 16 hours. Next, individual cell colonies were regrown in LB media at 37°C, collected at approximately OD_600_ =1 and stored at −80°C in LB media supplemented with 50% glycerol.

### Competition assay in a microbioreactor

Before the experiment, the milliliter-scale microbioreactor chips (the total volume of 2 mL) (**Fig. S5**) were γ-irradiated and sealed as part of the standard pre-inoculation sterile protocol^54^. The medium bottles and feed lines were autoclaved separately, and 0.22 µm-filters were installed between the bottles and the lines to prevent the microbioreactor from microbial contamination. Before each experiment, both the antibiotic-resistance populations and the sfGFP-positive non-resistant *E. coli* were grown separately for approximately 12 hours in the chemically defined media (New Minimal Media – NMM) containing 7.5 mM (NH_4_)_2_SO_4_, 8.5 mM NaCl, 22 mM KH_2_PO_4_, 50 mM K_2_HPO_4_, 1 mM MgSO_4_, 20 mM D-glucose, 50 mg/L of 20 canonical amino acids, 1 µg/mL each of Ca^2+^ and Fe^2+^, 0.01 µg/mL each of Cu^2+^, Zn^2+^, Mn^2+^, and Mo^2+^, 10 µg/mL of thiamine, and 10 µg/mL biotin) to achieve the exponential growth phase for each of the populations. Then, the cell cultures were mixed at approximately 1:1 ratio and immediately injected into a microbioreactor (the total volume of 2 mL) for the competition assay. Three microbioreactors were used simultaneously to support cell growth in (i) NMM media, (ii) NMM media supplemented with tavaborole (1µg/mL), and (iii) NMM media supplemented with norvaline (0.5 mM). Each of the microreactors was operating in the turbidostatic mode, maintaining the OD_600_ of mixed populations at 1, with the fermentation temperature was controlled at 37±0.1°C throughout the experiment. Each day, 100 µL of *E. coli* populations (5% of the total population size) were ejected from the each of the microbioreactors and used for flow cytometry analysis to immediately assess the ratio between sfGFP-positive and sfGFP-negative cells.

### Flow Cytometry

Analysis of live cells by flow cytometry was carried out on Attune NxT flow cytometer (Invitrogen). For each measurement, we used 100,000 cells. FlowJo v10 software was used to analyze the flow cytometry data.

## ACKNOWLEDGMENTS

We thank the following scientists for insightful discussions and perceptive comments on various drafts of this manuscript, and for bringing to our attention gaps in our knowledge and holes in our logic: Jessica Cunningham Reynolds (Moffitt Cancer Center), Alita Burmeister and Richard Prum (Department of Ecology and Evolutionary Biology, Yale), Jeffrey Tharp, Jon Fischer, Kazuaki Amikura, and Oscar Vargas-Rodriguez (the Department of Molecular Biophysics and Biochemistry, Yale), Antonia van den Elzen (Department Cellular & Molecular Physiology, Yale), Haissi Cui (The Scripps Research Institute), Javid Babak (Centre for Infectious Diseases at Tsinghua University School of Medicine), and Ya-Ming Hou and her laboratory members (Thomas Jefferson University). We also gratefully acknowledge the laboratory of Alanna Schepartz (currently at UC, Berkeley) for providing access to the flow cytometer, and Kenneth Nelson (the Yale Flow Cytometry Core Facility) for his technical assistance. This work was supported by the Guangdong Provincial Key Laboratory of Genome Read and Write (No. 2017B030301011 to Y.S.) and the National Institutes of Health Grant R35GM122560 (to D.S.).

## REFERENCES

1 Stankova, K. Resistance games. Nat Ecol Evol 3, 336–337, doi: 10.1038/s41559-018-0785-y (2019).

2 Cunningham, J. J. A call for integrated metastatic management. Nat Ecol Evol 3, 996–998, doi: 10.1038/s41559-019-0927-x (2019).

3 Thomas, F. et al. Is adaptive therapy natural? PLoS Biol 16, e2007066, doi: 10.1371/journal.pbio.2007066 (2018).

4 Hochberg, M. E. An ecosystem framework for understanding and treating disease. Evol Med Public Health 2018, 270–286, doi: 10.1093/emph/eoy032 (2018).

5 Zhang, J., Cunningham, J. J., Brown, J. S. & Gatenby, R. A. Integrating evolutionary dynamics into treatment of metastatic castrate-resistant prostate cancer. Nat Commun 8, 1816, doi: 10.1038/s41467-017-01968-5 (2017).

6 Cunningham, J. J., Gatenby, R. A. & Brown, J. S. Evolutionary dynamics in cancer therapy. Mol Pharm 8, 2094–2100, doi: 10.1021/mp2002279 (2011).

7 Maharjan, R. & Ferenci, T. The fitness costs and benefits of antibiotic resistance in drug-free microenvironments encountered in the human body. Environ Microbiol Rep 9, 635–641, doi: 10.1111/1758-2229.12564 (2017).

8 Melnyk, A. H., Wong, A. & Kassen, R. The fitness costs of antibiotic resistance mutations. Evol Appl 8, 273–283, doi: 10.1111/eva.12196 (2015).

9 Andersson, D. I. & Hughes, D. Antibiotic resistance and its cost: is it possible to reverse resistance? Nat Rev Microbiol 8, 260–271, doi: 10.1038/nrmicro2319 (2010).

10 Lenski, R. E. Bacterial evolution and the cost of antibiotic resistance. Int Microbiol 1, 265–270 (1998).

11 Spratt, B. G. Resistance to antibiotics mediated by target alterations. Science 264, 388–393, doi: 10.1126/science.8153626 (1994).

12 Munita, J. M. & Arias, C. A. Mechanisms of antibiotic resistance. Microbiol Spectr 4, doi: 10.1128/microbiolspec.VMBF-0016-2015 (2016).

13 Kim, S., Lieberman, T. D. & Kishony, R. Alternating antibiotic treatments constrain evolutionary paths to multidrug resistance. Proc Natl Acad Sci U S A 111, 14494–14499, doi: 10.1073/pnas.1409800111 (2014).

14 Nichol, D. et al. Steering evolution with sequential therapy to prevent the emergence of bacterial antibiotic resistance. PLoS Comput Biol 11, e1004493, doi: 10.1371/journal.pcbi.1004493 (2015).

15 Fuentes-Hernandez, A. et al. Using a sequential regimen to eliminate bacteria at sublethal antibiotic dosages. PLoS Biol 13, e1002104, doi: 10.1371/journal.pbio.1002104 (2015).

16 Zhao, B. et al. Exploiting temporal collateral sensitivity in tumor clonal evolution. Cell 165, 234–246, doi: 10.1016/j.cell.2016.01.045 (2016).

17 Mira, P. M. et al. Rational design of antibiotic treatment plans: a treatment strategy for managing evolution and reversing resistance. PLoS One 10, e0122283, doi: 10.1371/journal.pone.0122283 (2015).

18 Baker, S. J. et al. Discovery of a new boron-containing antifungal agent, 5-fluoro-1,3-dihydro-1-hydroxy-2,1-benzoxaborole (AN2690), for the potential treatment of onychomycosis. J Med Chem 49, 4447–4450, doi: 10.1021/jm0603724 (2006).

19 Rock, F. L. et al. An antifungal agent inhibits an aminoacyl-tRNA synthetase by trapping tRNA in the editing site. Science 316, 1759–1761, doi: 10.1126/science.1142189 (2007).

20 Zhang, P. & Ma, S. Recent development of leucyl-tRNA synthetase inhibitors as antimicrobial agents. Medchemcomm 10, 1329–1341, doi: 10.1039/c9md00139e (2019).

21 Jinna, S. & Finch, J. Spotlight on tavaborole for the treatment of onychomycosis. Drug Des Devel Ther 9, 6185–6190, doi: 10.2147/DDDT.S81944 (2015).

22 Manhas, R. et al. Leishmania donovani Parasites Are Inhibited by the benzoxaborole AN2690 targeting leucyl-tRNA synthetase. Antimicrob Agents Chemother 62, doi: 10.1128/AAC.00079-18 (2018).

23 Sonoiki, E. et al. Antimalarial benzoxaboroles target *Plasmodium falciparum* leucyl-tRNA synthetase. Antimicrob Agents Chemother 60, 4886–4895, doi: 10.1128/AAC.00820-16 (2016).

24 Palencia, A. et al. Cryptosporidium and *Toxoplasma* parasites are inhibited by a benzoxaborole targeting leucyl-tRNA synthetase. Antimicrob Agents Chemother 60, 5817–5827, doi: 10.1128/AAC.00873-16 (2016).

25 Xin, W. et al. Design and synthesis of alpha-phenoxy-N-sulfonylphenyl acetamides as Trypanosoma brucei Leucyl-tRNA synthetase inhibitors. Eur J Med Chem 185, 111827, doi: 10.1016/j.ejmech.2019.111827 (2020).

26 Zhang, F. et al. Discovery of N-(4-sulfamoylphenyl)thioureas as *Trypanosoma brucei* leucyl-tRNA synthetase inhibitors. Org Biomol Chem 11, 5310–5324, doi: 10.1039/c3ob40236c (2013).

27 Ding, D. et al. Discovery of novel benzoxaborole-based potent antitrypanosomal agents. ACS Med Chem Lett 1, 165–169, doi: 10.1021/ml100013s (2010).

28 Jacobs, R. T. et al. SCYX-7158, an orally-active benzoxaborole for the treatment of stage 2 human African trypanosomiasis. PLoS Negl Trop Dis 5, e1151, doi: 10.1371/journal.pntd.0001151 (2011).

29 Ding, D. et al. Design, synthesis, and structure-activity relationship of *Trypanosoma brucei* leucyl-tRNA synthetase inhibitors as antitrypanosomal agents. J Med Chem 54, 1276–1287, doi: 10.1021/jm101225g (2011).

30 Si, Y. et al. Antibacterial activity and mode of action of a sulfonamide-based class of oxaborole leucyl-tRNA-Synthetase inhibitors. ACS Infect Dis 5, 1231–1238, doi: 10.1021/acsinfecdis.9b00071 (2019).

31 Li, X. et al. Discovery of a potent and specific *M. tuberculosis* leucyl-tRNA synthetase Inhibitor: (S)-3-(Aminomethyl)-4-chloro-7-(2-hydroxyethoxy)benzo[c][1,2]oxaborol-1(3H)-ol (GSK656). J Med Chem 60, 8011–8026, doi: 10.1021/acs.jmedchem.7b00631 (2017).

32 Palencia, A. et al. Discovery of novel oral protein synthesis inhibitors of *Mycobacterium tuberculosis* that target leucyl-tRNA synthetase. Antimicrob Agents Chemother 60, 6271–6280, doi: 10.1128/AAC.01339-16 (2016).

33 Gudzera, O. I. et al. Identification of *Mycobacterium tuberculosis* leucyl-tRNA synthetase (LeuRS) inhibitors among the derivatives of 5-phenylamino-2H-[1,2,4]triazin-3-one. J Enzyme Inhib Med Chem 31, 201–207, doi: 10.1080/14756366.2016.1190712 (2016).

34 Gudzera, O. I. et al. Discovery of potent anti-tuberculosis agents targeting leucyl-tRNA synthetase. Bioorg Med Chem 24, 1023–1031, doi: 10.1016/j.bmc.2016.01.028 (2016).

35 Hu, Q. H. et al. Discovery of a potent benzoxaborole-based anti-pneumococcal agent targeting leucyl-tRNA synthetase. Sci Rep 3, 2475, doi: 10.1038/srep02475 (2013).

36 Palencia, A. et al. Structural dynamics of the aminoacylation and proofreading functional cycle of bacterial leucyl-tRNA synthetase. Nat Struct Mol Biol 19, 677–684, doi: 10.1038/nsmb.2317 (2012).

37 Zhao, H. et al. Analysis of the resistance mechanism of a benzoxaborole inhibitor reveals insight into the leucyl-tRNA synthetase editing mechanism. ACS Chem Biol 10, 2277–2285, doi: 10.1021/acschembio.5b00291 (2015).

38 Sarkar, J., Mao, W., Lincecum, T. L., Jr., Alley, M. R. & Martinis, S. A. Characterization of benzoxaborole-based antifungal resistance mutations demonstrates that editing depends on electrostatic stabilization of the leucyl-tRNA synthetase editing cap. FEBS Lett 585, 2986–2991, doi: 10.1016/j.febslet.2011.08.010 (2011).

39 Gupta, A. et al. A Polymorphism in leuS confers reduced susceptibility to GSK2251052 in a clinical isolate of *Staphylococcus aureus*. Antimicrob Agents Chemother 60, 3219–3221, doi: 10.1128/AAC.02940-15 (2016).

40 O’Dwyer, K. et al. Bacterial resistance to leucyl-tRNA synthetase inhibitor GSK2251052 develops during treatment of complicated urinary tract infections. Antimicrob Agents Chemother 59, 289–298, doi: 10.1128/AAC.03774-14 (2015).

41 Li, L. et al. Naturally occurring aminoacyl-tRNA synthetases editing-domain mutations that cause mistranslation in *Mycoplasma* parasites. Proc Natl Acad Sci U S A 108, 9378–9383, doi: 10.1073/pnas.1016460108 (2011).

42 Melnikov, S. V. et al. Error-prone protein synthesis in parasites with the smallest eukaryotic genome. Proc Natl Acad Sci U S A 115, E6245–E6253, doi: 10.1073/pnas.1803208115 (2018).

43 Karkhanis, V. A., Mascarenhas, A. P. & Martinis, S. A. Amino acid toxicities of Escherichia coli that are prevented by leucyl-tRNA synthetase amino acid editing. J Bacteriol 189, 8765–8768, doi: 10.1128/JB.01215-07 (2007).

44 Cvetesic, N. et al. Proteome-wide measurement of non-canonical bacterial mistranslation by quantitative mass spectrometry of protein modifications. Sci Rep 6, 28631, doi: 10.1038/srep28631 (2016).

45 Ribas de Pouplana, L., Santos, M. A., Zhu, J. H., Farabaugh, P. J. & Javid, B. Protein mistranslation: friend or foe? Trends Biochem Sci 39, 355–362, doi: 10.1016/j.tibs.2014.06.002 (2014).

46 Bacevic, K. et al. Spatial competition constrains resistance to targeted cancer therapy. Nat Commun 8, 1995, doi: 10.1038/s41467-017-01516-1 (2017).

47 Elsa Hansen, J. K., Robert J. Woods, Andrew F. Read, Kevin B. Wood. Antibiotics can be used to contain drug-resistant bacteria by maintaining sufficiently large sensitive populations. bioRxiv, doi: https://doi.org/10.1101/638924 (2019).

48 Hansen, E., Woods, R. J. & Read, A. F. How to use a chemotherapeutic agent when resistance to it threatens the patient. PLoS Biol 15, e2001110, doi: 10.1371/journal.pbio.2001110 (2017).

49 Sharan, S. K., Thomason, L. C., Kuznetsov, S. G. & Court, D. L. Recombineering: a homologous recombination-based method of genetic engineering. Nat Protoc 4, 206–223, doi: 10.1038/nprot.2008.227 (2009).

50 Li, H. & Durbin, R. Fast and accurate short read alignment with Burrows-Wheeler transform. Bioinformatics 25, 1754–1760, doi: 10.1093/bioinformatics/btp324 (2009).

51 McKenna, A. et al. The Genome Analysis Toolkit: a MapReduce framework for analyzing next-generation DNA sequencing data. Genome Res 20, 1297–1303, doi: 10.1101/gr.107524.110 (2010).

52 Francklyn, C. S., First, E. A., Perona, J. J. & Hou, Y. M. Methods for kinetic and thermodynamic analysis of aminoacyl-tRNA synthetases. Methods 44, 100–118, doi: 10.1016/j.ymeth.2007.09.007 (2008).

53 Splan, K. E., Musier-Forsyth, K., Boniecki, M. T. & Martinis, S. A. *In vitro* assays for the determination of aminoacyl-tRNA synthetase editing activity. Methods 44, 119–128, doi: 10.1016/j.ymeth.2007.10.009 (2008).

54 Perez-Pinera, P. et al. Synthetic biology and microbioreactor platforms for programmable production of biologics at the point-of-care. Nat Commun 7, 12211, doi: 10.1038/ncomms12211 (2016).

