## Supplementary Data 2 for "Exploiting evolutionary trade-offs to combat antibiotic resistance"

**Supplementary Data 2 | Time-resolved genome sequencing of the evolving populations A and B.** The sequencing data were then deposited to the CNGB Nucleotide Sequence Archive, CNSA: <https://db.cngb.org/cnsa>. The accession number is CNP0000842.
