## Supplementary Information for "Exploiting evolutionary trade-offs to combat antibiotic resistance"

**Tables**

Table S1. Mutations observed in the tavaborole-resistant *E. coli* colonies.

Table S2. Primers used in the study.

**Figures**

Fig. S1 | Tavaborole-resistance evolution in *E. coli*.

Fig. S2 | Mutations in the *leuS* gene confer tavaborole-resistance.

Fig. S3 | Time-resolved whole-population genome sequencing illustrates rapid propagation of *leuS* mutations in the evolving *E. coli*.

Fig. S4 | Mutations in LeuRS editing site appear to impair tRNA binding to the editing domain.

Fig. S5 | The principal component of the microbioreactor used for the competition evolutionary experiment.

**Data:**

Supplementary Data 1 | Sanger sequencing of 120 colonies from the evolved tavaborole-resistant populations of *E. coli*.

Supplementary Data 2 | Time-resolved genome sequencing of the evolving populations A and B. (the file contains links to the sequencing databases)

Supplementary Data 3 | The LeuRS-coding plasmid *leuS*-pBAD28.

Supplementary Data 4 | The integration plasmid to insert sfGFP-coding gene into the *E. coli* genome.

| Population A | Population B | Population C | Population D | Population E | Population F |
|---|---|---|---|---|---|
| R344S | R344C | G225E | M336MAV | V338D | G331C |
| R344S | R344C | R344S | M336MAV | V338D | G229V G331C |
| R344S | R344S | G225E | M336MAV | V338D | G225E G331R |
| G225E | R344S | R344S | M336MAV | R344S | G229V |
| G225E | R344S | R344S | M336MAV | V338D | G331C |
| G225E | R344C | G225E | M336MAV | V338D | G331C |
| R344S | R344S | G225E | M336MAV | V338D | G225E G331C |
| G225E | Y330F | G225E | M336MAV | V338D | G225E |
| R344S | R344S | R344S | Q269P | R344S | G331C |
| G225E | Y330F | A334E | M336MAV | R344S | G225E G331C |
| G225E | R344C | G229V | M336MAV | R344S | G225E |
| R344S | R344S | R344S | M336MAV | V338D | G331C |
| R344S | L354R | A334E | Q269P | R344S | G331C |
| G225E | R344S | R344S | M336MAV | V338D | G225E |
| G229V | G229T | M336I | Q269P | V338D | G225E |
| G229V | R344C | G225E | M336MAV | R344S | G331C |
| G225E | R344S | G225E | M336MAV | V338D | G331C |
| G225E | R344S | R344S | M336MAV | V338D | G229V |
| G225E | R344S | R344S | M336MAV | V338D | G331C |
| G225E | V335VMA | R344S | M336MAV | V338D | G225E |

**Table S1. Mutations observed in the tavaborole-resistant *E. coli* colonies.** The table summarizes mutations that were observed in *leuS* gene in 120 tavaborole-resistant colonies of the evolved *E. coli*. The corresponding Sanger sequencing data are deposited as (**Supplementary Data 1**).

| Primer | Sequence | Description |
| --- | --- | --- |
| **1** | CATCCGCCAAAACAGCTTAGCCAACGACCAGATTGAGGAGTTT | *leuS* amplification |
| **2** | AGCAGCGGCCAAGAGCAATACCGC | *leuS* amplification |
| **3** | TGGATAAACTGGATCACTGGCCAGAC | editing domain segment amplification |
| **4** | TTACCACATCTTCCGGCAGGATCAC | editing domain segment amplification |
| **5** | GCTGTTTTGGCGGATGAGAGAAG | pBAD28 amplification for *leuS* cloning |
| **6** | GCTAGCCCAAAAAAACGGGTATGG | pBAD28 amplification for *leuS* cloning |
| **7** | GGTGATCCTGCCGGAAGATGTGGTAA | pBAD28 amplification for ed-domain cloning |
| **8** | GGTGATCCTGCCGGAAGATGTGGTAA | pBAD28 amplification for ed-domain cloning |
| **9** | TTCTGTTTTATCAGACCGCTTCTGCG | pBAD28 insert analysis |
| **10** | ATAGCATTTTTATCCATAAGATTAGCGGATCC | pBAD28 insert analysis |
| **11** | CCTAATACGACTCACTATAGCCGAAGTGGCGAAATCGGTAGA | DNA template for $tRNA^{Leu}$ synthesis |
| **12** | GCCGAAGTGGCGAAATCGGTAGACGCAGTTGATTCAAAATCAACCGTAGAAATACGTGC | DNA template for $tRNA^{Leu}$ synthesis |
| **13** | TGGTGCCGAAGGCCGGACTCGAACCGGCACGTATTTCTACGGTTGATTTTGAATCAAC | DNA template for $tRNA^{Leu}$ synthesis |

**Table S2. Primers used in the study.**

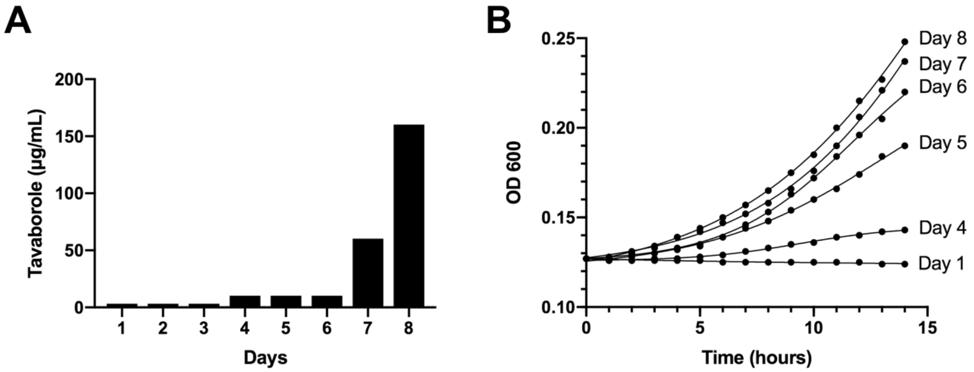


**Fig. S1 | Tavaborole-resistance evolution in *E. coli*. A.** The diagram shows tavaborole concentrations that were used to evolve resistance in 10 populations of *E. coli* that were grown in parallel. Tavaborole was added to the growth media to 2.5 ug/ml concentration. As the populations showed the sign of tavaborole-resistance, tavaborole concentrations were gradually increased to the final concentration of 160 ug/ml. **B.** Growth curves of one of the evolving E. coli culture (lineage 1) collected at days 1-8 of the experiment and regrown simultaneously in the presence of tavaborole (16ug/ml). As the diagram shows, the initial cell population (collected at Day 1) cannot grow in the presence of tavaborole (16ug/ml), however in the course of the evolution experiment the population acquires the ascendingly rapid growth in the presence of the drug.

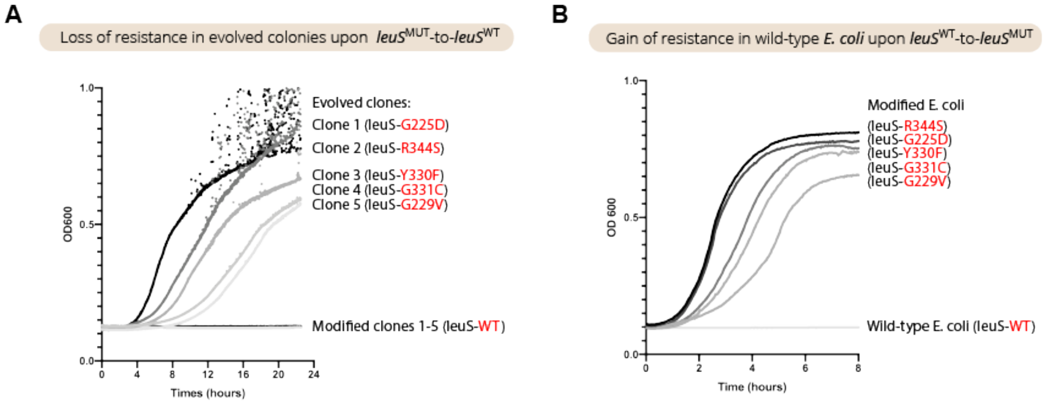


**Fig. S2 | Mutations in the *leuS* gene confer tavaborole-resistance.** The panels show growth rate assays of the evolved and genetically engineered strains of *E. coli* to test if mutations in the editing domain of LeuRS indeed confer tavaborole resistance. **A.** Growth curves comparing two sets of *E. coli* cells: the evolved *E. coli* clones (clones 1–5), in which *leuS* contained one of the five most frequently observed mutations, and the same clones after their *leuS* gene had been replaced with the wild-type *leuS* gene. When the evolved clones were modified and their *leuS* sequence was reverted to the wild-type, they lost their tavaborole resistance, indicating that the tavaborole-resistant phenotype is determined by mutations in the *leuS* gene. **B.** Growth curves comparing wild-type *E. coli* with the derived clones in which the *leuS* gene was modified to introduce one of the five most frequently observed mutations in the *leuS* gene. When *E. coli* acquire one mutation in *leuS* in their genomic DNA, they become resistant to tavaborole.

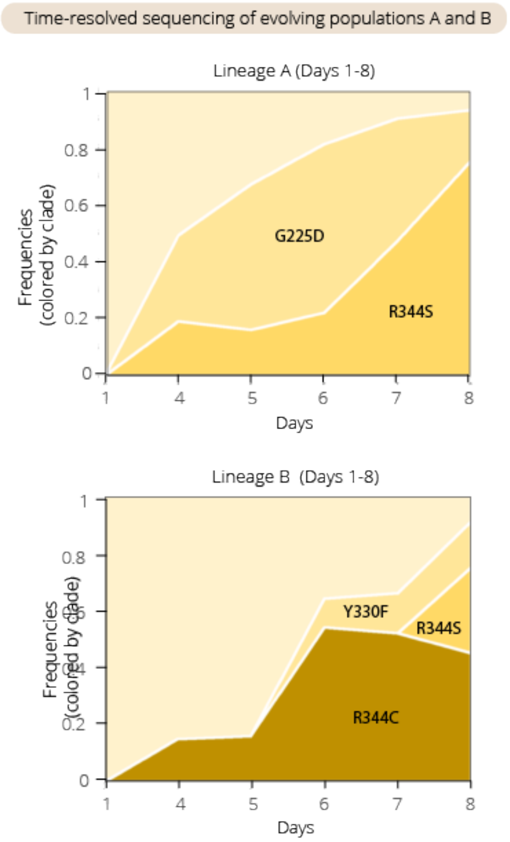


**Fig. S3 | Time-resolved whole-population genome sequencing illustrates rapid propagation of *leuS* mutations in the evolving *E. coli*.** Time-resolved whole-population DNA sequencing for two independent populations of *E. coli* (populations A and B) growing in the presence of tavaborole. The panels show that mutations in the *leuS* gene accumulated in a time-dependent manner and were eventually present in the majority of the evolving cells, illustrating that the majority of tavaborole-resistant *E. coli* have mutated editing domain in LeuRS.

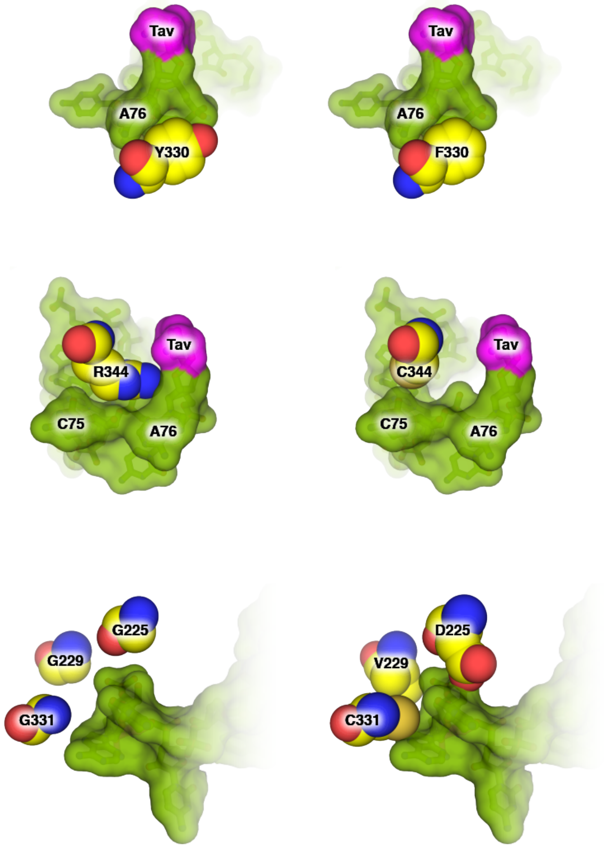


**Fig. S4 | Mutations in LeuRS editing site appear to compromise tRNA binding to the editing domain.** The panels show fragments of the crystal structure of bacterial LeuRS/tRNA/tavaborole complex (left panels, pdb id **2v0g**) and the hypothetic structures of LeuRS mutants (right panels) to illustrate an apparent mechanism by which LeuRS mutations prevent tRNA accommodation in the editing site. The structure suggests that several mutations confer tavaborole-resistance by disrupting tRNA contacts with LeuRS editing domain (such as the hydrogen bonds between Y330 residue and the 3′-terminal phosphate in $tRNA^{Leu}$ , or the salt bridging-contact between R344 and the 3′-terminal phosphate in $tRNA^{Leu}$). Other mutations, including G225D, G229V, and G331C, appear to create a steric clash between $tRNA^{Leu}$ and LeuRS, thereby preventing $tRNA^{Leu}$ binding to the editing site.

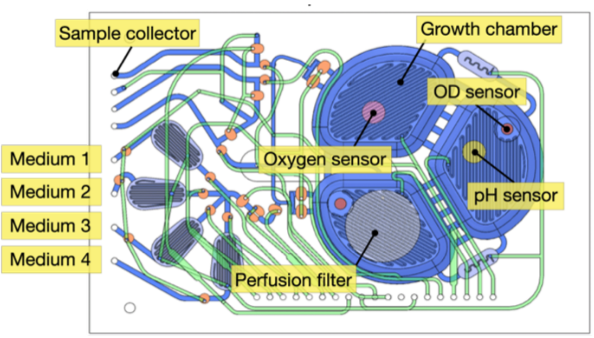


**Fig. S5 | The principal component of the microbioreactor used for the competition evolutionary experiment.** The scheme shows the milliliter-scale microbioreactors (as described in Ref. 49) used for the *E. coli* competition assay. Each reactor is a polycarbonate-PDMS membrane-polycarbonate sandwiched chip with active microfluidic circuits that are equipped for pneumatic routing of reagents, precise peristaltic injections, growth chamber mixing, and fluid extraction. Each chip has a total volume of 2 mL and allows continuous growth of a cell culture in a turbidostatic mode.
